# Improved Scalar Deregression of Estimated Breeding Values

**DOI:** 10.64898/2026.10.12.758488

**Authors:** Norbert Reinsch

## Abstract

**Background:** In genomic studies of animals and plants, analyses are often split into two stages. In the first stage, the average phenotype of each genotyped candidate is calculated from its estimated breeding value (EBV). This average is adjusted for all other effects and is known as the deregressed breeding value. Scalar deregression removes the inherent shrinkage from the original EBVs candidate-by-candidate, rather than for a vector of EBVs. A previously published standard method is easy to apply but implicitly assumes no inbreeding.

**Results:** An improved method uses each candidate’s and its parents’ inbreeding coefficients and the three-by-three prediction error covariance matrix of their EBVs. The candidate’s effective number of observations can be derived as the root of a second-order polynomial, and the deregressed EBV is then obtained. Two mathematically equivalent polynomials differed in numerical stability.

Applications to small examples demonstrated that the improved method produced deregressed EBVs unbiased by inbreeding, whereas the standard method underestimated their absolute magnitude. The same prediction error covariance matrix also allows filtering out uninformative candidates if their estimated Mendelian sampling term has zero reliability.

**Conclusions:** The improved method requires access to selected off-diagonal elements of the inverted coefficient matrix of the mixed model equations. The resulting deregressed EBVs and their appropriate weights are unbiased by inbreeding, thereby improving the baseline for the second stage of the analysis. Additionally, uninformative observations can be reliably filtered out. Studies involving traits with low heritability or sparsely recorded phenotypes are expected to benefit most.

## Background

In animal and plant breeding, genome-wide association studies (GWAS) are frequently performed in two stages. In the first stage, a mixed model adjusts all phenotypes for systematic environmental effects, accounts for covariance among observations by possibly including a range of non-genetic random effects, and provides a best linear unbiased predictor (BLUP) of the genetic merit of all animals with a phenotype and their relatives. These estimated breeding values (EBVs) summarize all known phenotypic information about genotyped candidates in a single quantity. There is, however, a broad consensus that estimated breeding values of genotyped candidates should not be carried into the second stage of an association study (Garrick et al., 2009; Ekine et al., 2014; Holland and Piepho, 2024; Verbyla, 2023; and the references therein). The main underlying reason is that BLUPs are shrunken towards their expectation, either the family means or the population mean, the extent of shrinkage depending on the amount of available information, which varies with the number of observations and the trait’s heritability. This has led to the development of deregressed EBVs (dEBVs) as pseudo-phenotypes. These eliminate shrinkage from the original EBVs, while providing the desired summary of the average available phenotypic information per genotyped candidate. Passing dEBVs and their appropriate weights on to the second stage of an analysis avoids reduced detection power, an increased false-positive rate, and biased QTL effect sizes, all of which are associated with using plain EBVs (Ekine et al., 2014).

Deregressing single EBVs separately is known as scalar deregression, as opposed to processing the full vector of EBVs (Banos et al., 1993; Jairath et al., 1998) in one go, which is called matrix deregression. A previously published scalar deregression method (Garrick et al., 2009) has widely become standard. For each particular candidate, it relies on its EBV, the EBVs of its parents, and their respective reliabilities. Each candidate’s resulting dEBV represents the information that is provided by the phenotype(s) of the candidate and its offspring. Contributions from ancestral phenotypes and sibs are removed with the help of the average EBV of the candidate’s parents.

Often, reliabilities for use in the standard method can be calculated from the diagonal elements of the inverse coefficient matrix of the mixed model equations. Relevant statistical packages for genetic analyses provide this information, typically as prediction error standard deviations. A practical advantage of the standard method is that it remains applicable when reliabilities are only available as approximations from large-scale systems for genetic evaluation, where EBVs are calculated iteratively without using the inverse coefficient matrix.

The scalar deregression method of Garrick et al. (2009) implicitly assumes that each candidate’s parents are unrelated and that neither the candidate nor its parents are inbred. Clearly, that assumption will frequently be violated in livestock data and in experimental genetics. Further, when researchers applied scalar deregression to traits with low heritability, they have frequently filtered out meaningless dEBVs by demanding that their weights surpass a minimum threshold (e.g., Sevillano et al. 2015; Gertz et al., 2016). This practice is especially relevant for traits related to fertility and health. However, filtering out pseudo-phenotypes with a low weight introduces a certain degree of arbitrariness, as inaccurate but valid information may be lost, while, at the same time, the weight of actually meaningless dEBVs may pass the threshold.

This manuscript considers situations where the coefficient matrix of the mixed model equations can be inverted and a software package like ASReml (Gilmour et al., 2015) provides access not only to the diagonal of the inverse coefficient matrix, but also to selected off-diagonal elements. For our pursuit, we rely on the three-by-three prediction error covariance matrix of the EBVs of each candidate and its parents. A modified scalar deregression procedure is derived that capitalizes on these prediction error covariances. These covariances also allow for the computation of the reliability of the candidate’s estimated Mendelian sampling term, which is proposed as a criterion for filtering out observations with no information content beyond the family mean. The parent average is removed similar as in the method of Garrick et al. (2009). The improved approach fully accounts for inbreeding and relationships between parents and the genotyped candidate. Toy examples demonstrate how the method works in various situations, and simulated phenotypes for two of these examples allow for a comparison of dEBVs from the traditional and the new approach.

## Methods

### Theory

We assume a population of pedigreed individuals with phenotypic data for a trait of interest. A mixed model comprising additive genetic effects can then be fitted to these data. Our interest is in de-regressing the EBVs of a subgroup of all individuals, which are considered to be genotyped. The resulting deregressed EBVs, together with their weights, will later serve as dependent variables in a GWAS or other type of second-stage analysis.

#### Relationships and inverse relationships

For each candidate three inbreeding coefficients *F*_*i*_, *F*_*s*_, and *F*_*d*_, can be retrieved from the pedigree, whereby indices indicate the candidate (i), its sire (s) and its dam (d). Then, the tree-by-three additive-genetic relationship matrix **A**_*sdi*_ of such a trio can be expressed in terms of their inbreeding coefficients as:

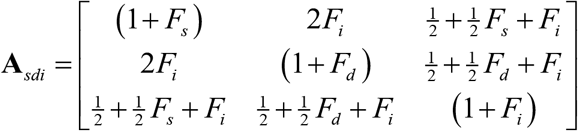

Therefrom, we get the diagonal of the two-by-two relationship matrix **A**_*pm*_ of the average breeding value of the parents *p*_*a*_ and the Mendelian sampling deviation *m* of the candidate.

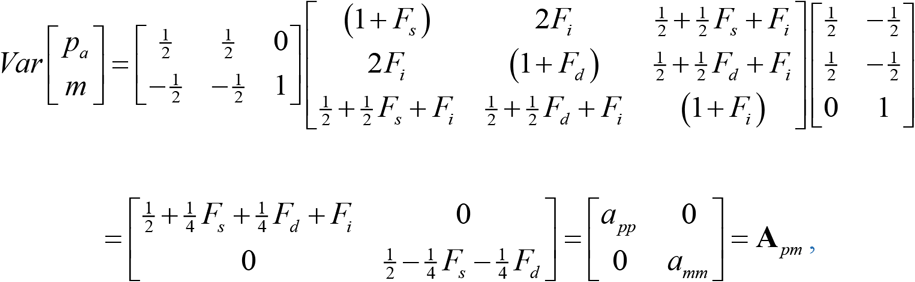

with diagonal elements *a*_*pp*_ and *a*_*mm*_ .

Likewise, the relationship between the average breeding value of the parents *p*_*a*_ and the breeding value *b* of the candidate is:

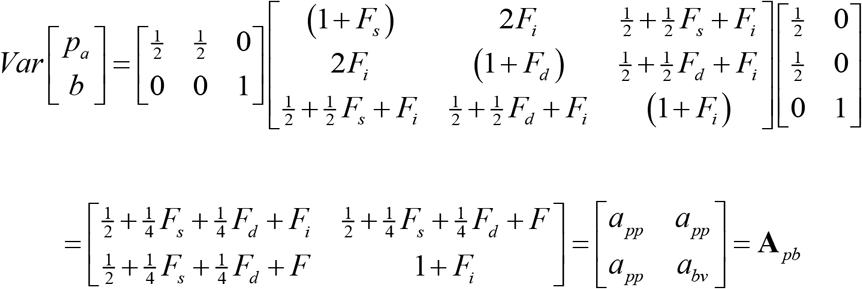

In the following we will make use of the inverse of the latter relationship matrix.

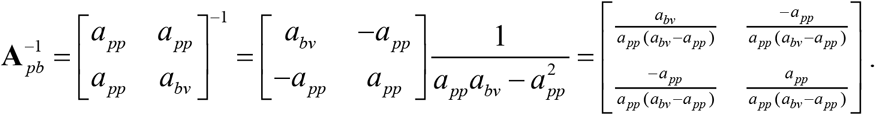

Using the elements in **A**_*pb*_ we can express the variance of the candidate’s Mendelian sampling effect as *a*_*pp*_ + *a*_*bv*_ − 2*a*_*pp*_ = *a*_*bv*_ − *a*_*pp*_ = *a*_*mm*_ . By using that result, the inverse can also be written as

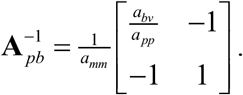

### Effective number of observations

As a first step of the deregression procedure, the unknown effective number of observations 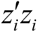 that is behind the precision of the candidate’s EBV has to be determined. To this end, we make use of the three-by-three prediction error covariance matrix of the EBVs of the candidate and its parents and transform it into the two-by-two prediction error matrix of the parent average and the candidate’s EBV.

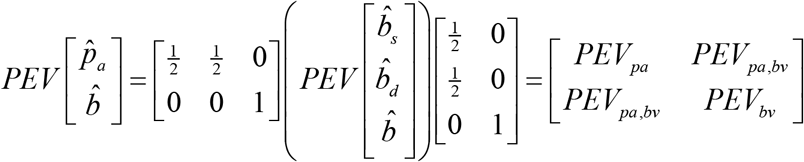

The two diagonal elements of the latter matrix, *PEV*_*pa*_ and *PEV*_*bv*_ provide us with the necessary information on the precision of the parent average and the candidate’s EBV, allowing for the computation of the effective number of observations.

Thereby, we follow the previously published approach of Garrick et al. (2009) in so far as we depart from two BLUP-like equations involving the known estimates for the parent average 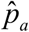 and the candidate’s breeding value 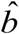 as random effects. However, here we account for inbreeding by employing the relationship matrix **A**_*pb*_ from above in setting up the coefficient matrix. The right-hand side can only later be determined, once the effective numbers of observations have been determined. Then, the resulting equations are

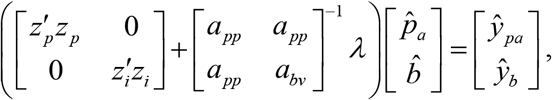

Whereby 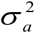 is the additive-genetic variance, 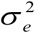 is the residual variance and 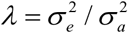. Equating the diagonal elements of the inverse coefficient matrix to the respective prediction error variances results in two equations that involve the effective number of observations 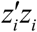 for the candidate and its counterpart 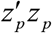 for the parent average.

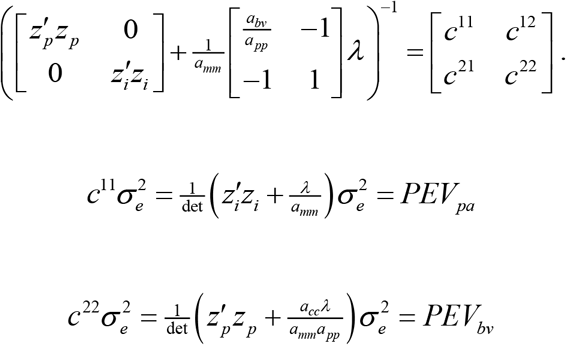

The relationship between 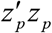 and 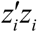 is linear, as can be found by dividing one of the two equations by the other. This relationship can be summarized as

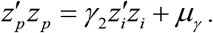

That can be used to replace 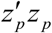 in one of the two original equations and leads to a second-order polynomial in 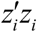

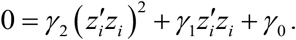

All coefficients of the latter linear and quadratic equations can be expressed in terms of prediction error variances, relationships, the variance ratio and the residual variance as

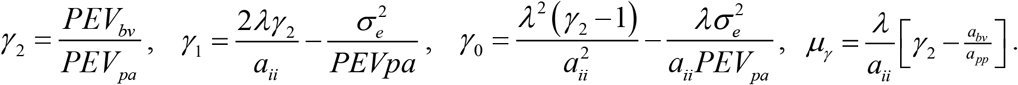

Then the solution to the quadratic equation is

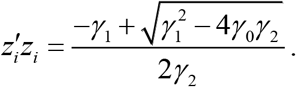

Alternatively, the set{*γ*_2_, *γ*_1_,*γ*_0_, *μ*_*γ*_ }could be expressed as functions of reliabilities instead of PEVs. That does not affect the resulting values whatsoever.

#### Deregressed EBV and reliability

After the efficient number of observations have been determined, the now fully known coefficient matrix can be multiplied with the EBVs from the BLUP evaluation to reconstruct the right-hand side of the two equations. In particular, we are interested in its second element *ŷ*_*b*_, which can be understood as the sum of all observations for the candidate, adjusted for all other effects in the model. Observations from the candidate’s offspring also contribute to this quantity. Finally, dividing this sum by the effective number of observations turns it into an average, which can serve as the dEBV or pseudo-phenotype.

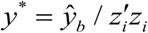

The reliability of the dEBV is solely a function of the effective number of observations for the candidate, the variance ratio, and the candidate’s inbreeding coefficient:

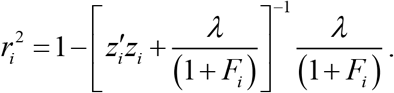

The effect of parental phenotypes, and phenotypes of other relatives contributing to the PA, like sibs, on the reliability is not taken into account (is “removed”, Garrick et al., 2009). This is the reason why the reliability of any dEBV never exceeds the reliability of its corresponding EBV.

Finally, the reliability of the dEBV determines its weight in a second stage of analysis:

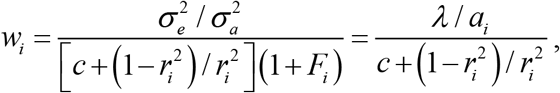

where the coefficient c is the proportion of genetic variance not explained by markers. If the inbreeding coefficient *F*_*i*_ of the candidate is zero, the weight equals that given in Garrick et al. (2009).

#### Alternative equations for the effective number of observations

Alternatively, we may transform the vector of genetic effects into the parent average and the candidate’s

Mendelian sampling term with the help of a transformation matrix **T**

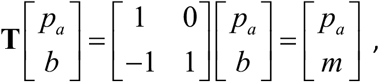

where the reverse transformation is

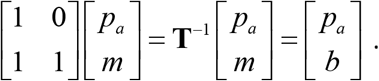

Then, the underlying incidence matrix **Z** accordingly becomes

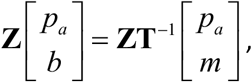

and consequently, we get

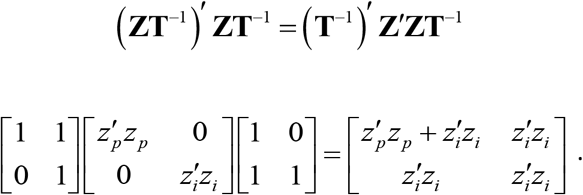

Finally, we obtain the following transformed equations as a starting point for our further derivations:

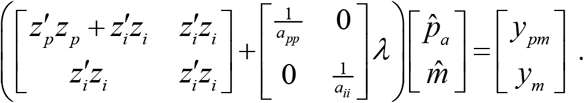

Note that the right-hand side of these equations is also a transformed version of that shown above (see also below). We denote the inverse of the coefficient matrix as

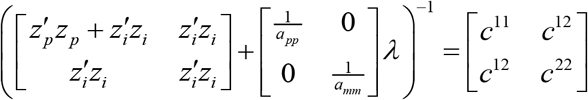

and relate the first diagonal of this inverse to the reliability 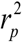 of the estimated parent average and the second diagonal element to the reliability 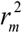 of the candidate’s estimated Mendelian sampling effect.

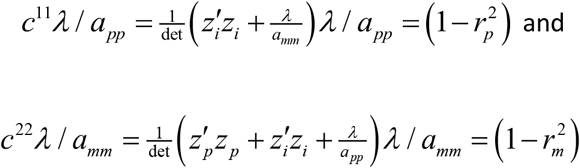

Thereby det is the determinant of the coefficient matrix.

A linear equation converting 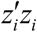 into 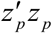 can be derived by dividing the first equation by the second and solving for 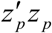 .

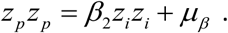

Substituting that equality into the first equation yields an alternative second-order polynomial in the unknown 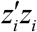 .

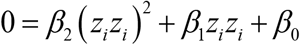

All four coefficients for these two equations are functions of the variance ratio *λ*, the reliabilities 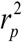 and 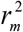, and the pedigree-derived quantities *a*_*pp*_ and *a*_*mm*_ :

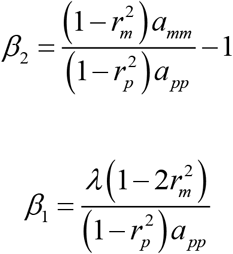

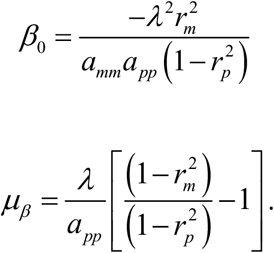

Note that expressing the latter four coefficients in terms of PEVs rather than reliabilities leaves the resulting values unaltered and yields the same polynomial, just as with the first polynomial above. With all elements of the coefficient matrix known, we obtain the right-hand side of the equations by multiplying the coefficient matrix by the vector of estimates, as in the previous version. Our interest, however, is in the right-hand side of two equations involving the average EBV 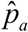 of the parents and the candidate’s EBV 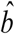, not its estimated Mendelian sampling term. The following back-transformation defines the corresponding right-hand side:

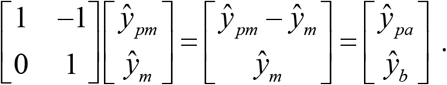

The transformation formula shows that the second element remains untouched, and 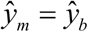 is the accumulated phenotypic information available for the estimation of the candidate’s EBV, except for all information captured in the average parental EBV. Taking the average gives the identical deregressed EBV

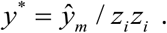

Finally, the reliability of that EBV and the corresponding weight remain the same as previously described The set {*β*_2_, *β*_1_, *β*_0_, *μ*_*β*_ } of coefficients leads to a quadratic equation that has *β*_2_ in the denominator. In case the prediction error variance *PEV*_*mm*_ of the Mendelian sampling effect and that of the parent average are equal, that coefficient becomes zero, and the quadratic equation has a discontinuity:

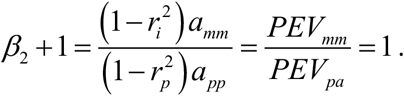

The solution for 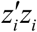 in such a situation is given by the rule of L’Hospital.

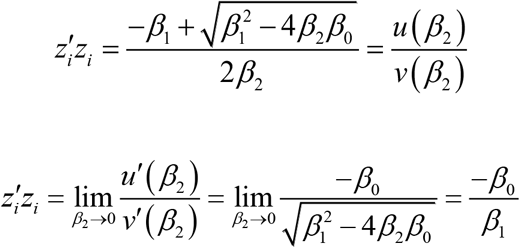

and

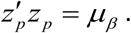

In practice such a discontinuity does not occur in the quadratic equation that relies on{*γ*_2_, *γ*_1_,*γ*_0_, *μ*_*γ*_ }. The reason is that it has the ratio *γ*_2_ = *PEV*_*bv*_ / *PEV*_*pa*_ in the denominator, which is always positive in relevant applications with non-zero prediction-error variances.

#### Lack of information

The quadratic formula in terms of {*γ*_2_, *γ*_1_,*γ*_0_, *μ*_*γ*_ }makes the important case obvious where the reliability 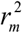 of a candidate’s the Mendelian sampling term is zero. Then the coefficient *β* also becomes zero and the quadratic equation delivers 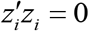. This propagates to a zero reliability of the deregressed EBV of *r*^2^ = 0, which itself has no defined value due to division by zero. As a consequence, the EBVs of such candidates cannot be deregressed. Instead, they have to be filtered out from any second stage analysis due to a lack of information. Ideally, one first computes 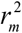 from the PEV-matrix for each candidate. The candidate’s EBV is only deregressed when 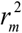 exceeds a small operational value such as *ε* = 0.02, otherwise it is filtered out.

### Illustrative Examples

#### Specification of examples

The estimation of the effective number of observations *z*_*i*_ *z*_*i*_ for a candidate is illustrated through six small examples based on a simple pedigree involving parents which are related but not inbred. The relationship between them was set to 0.5, resulting in the candidate’s inbreeding coefficient of 0.25. Additionally, two offspring of the candidate, with no known other parent, are included in the relationship matrix **A** . Prediction error variances and covariances were calculated by inverting the coefficient matrix of a model that considers only the additive-genetic effect as its single random effect and includes no fixed effects at all.

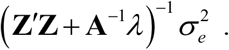

The genetic variance was assumed as one in all examples. The incidence matrix **Z** was varied to match the specifications of each of the examples as given in **Table 1**. More details, such as the numerator-relationship matrix and the design matrices, are displayed in the **Appendix**.

**Table 1:** Specification of six illustrative examples used for scalar deregression. They all rely on the same pedigree but differ in the numbers of observations (np, nc, no: number of observations provided by the parents, the candidate, and its offspring), and the variance ratio *λ* . The resulting prediction error variances for the parent average (PEVp), the candidate’s Mendelian sampling effect (PEVi), and the candidate’s estimated breeding value (PEVbv) are followed by the respective reliabilities in the same order.

| Ex. | number of observations | | | $\lambda$ | PEVs | | | reliabilities | | |
| --- | --- | --- | --- | --- | --- | --- | --- | --- | --- | --- |
| | np | nc | no | | PEV<sub>p</sub> | PEV<sub>i</sub> | PEV<sub>bv</sub> | $r_p^2$ | $r_i^2$ | $r_{bv}^2$ |
| 1 | 2 | 5 | 0 | 4 | 0.38424 | 0.36453 | 0.45320 | 0.48768 | 0.27094 | 0.63744 |
| 2 | 2 | 5 | 0 | 3 | 0.34375 | 0.34375 | 0.37500 | 0.54167 | 0.31250 | 0.70000 |
| 3 | 2 | 0 | 2 | 4 | 0.66038 | 0.47601 | 0.94057 | 0.31080 | 0.04798 | 0.24755 |
| 4 | 2 | 1 | 2 | 4 | 0.46816 | 0.43505 | 0.76151 | 0.37579 | 0.12990 | 0.39080 |
| 5 | 2 | 0 | 0 | 4 | 0.54545 | 0.50000 | 1.04545 | 0.27273 | zero | 0.16364 |
| 6 | 2 | 0 | 0 | 3 | 0.50000 | 0.50000 | 1.00000 | 0.33333 | zero | 0.20000 |

In the first example ( *λ* = 4 ), the candidate had five observations, and no observations were contributed by offspring. Each of the parents had a single phenotypic record, and this was the same in all other examples. The second example is identical to the first, except for a variance ratio of *λ* = 3. In the third and fourth examples the variance ratio was again *λ* = 4 and both offspring each provided a single phenotypic record, while the candidate did so only in example four. Finally, in examples five ( *λ* = 4 ) and six ( *λ* = 3) neither the candidate nor its offspring had any phenotypic information. Table 1 also presents the resulting prediction error variances for the parent average, the candidate’s Mendelian sampling term, and its EBV, together with the corresponding reliabilities.

In a simulation, two hundred observation vectors were generated for each of the examples two and three. Therefrom the candidate’s dEBV was then calculated twice for each repetition, using the new and the traditional method. Resulting dEBVs from both methods were compared by regressing old on new dEBVs.

## Results

The newly derived method correctly reproduced the number of “effective” observations for the candidate in all examples (**Table 2**, column six). The value of 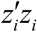 for the candidate was reproduced with no error as five in the first and second examples. In example three, with no phenotype of the candidate but two offspring with a phenotypic record, the result was 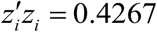, as expected.

**Table 2.** Results are shown for the newly derived scalar deregression method (new) and a previously published approach (old), when they were applied to all examples. In the old method the reliability of the parent average is 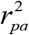 . For both versions, the effective number of observations for the candidate 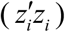 and for the parent average 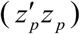 and are given, along with the resulting reliability of the candidate’s deregressed breeding value (r^2^).

| example | $r_{pa}^2$ | $z'_p z_p$ | | $z'_i z_i$ | | $r^2$ | |
| --- | --- | --- | --- | --- | --- | --- | --- |
|  | old | old | new | old | new | old | new |
| 1 | 0.19677 | 1.55129 | 2.0000 | 6.67907 | 5.0000 | 0.63 | 0.61 |
| 2 | 0.22098 | 1.59844 | 2.0000 | 6.64736 | 5.0000 | 0.69 | 0.66 |
| 3 | 0.13044 | 2.04493 | 2.0000 | 0.86265 | 0.42667 | 0.22 | 0.12 |
| 4 | 0.15481 | 1.89837 | 2.0000 | 2.14169 | 1.42667 | 0.35 | 0.31 |
| 5 | 0.11616 | 2.11797 | 2.0000 | 0.31501 | zero | 0.07 | zero |
| 6 | 0.14286 | 2.11363 | 2.0000 | 0.30073 | zero | 0.09 | zero |

This is two times 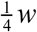, where *w* is the weight for each offspring’s phenotype in a reduced animal model (see the Appendix). Likewise, in example four 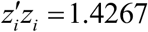, which is as in the third example but increased by one, because of the additional phenotype of the candidate. Finally, examples five and six correctly delivered zero effective observations, due to a zero reliability of the candidate’s estimated Mendelian sampling term. The number of effective observations for the parent average 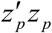 was correctly estimated as two in all examples. That value would be smaller if two observations had been assigned to one parent and none to the other, depending on the variance ratio.

For comparison, Table 2 also shows the outcomes of applying the method of Garrick et al. (2009) to all six examples. Note that the reliability of the parent average (**Table2**, first column) for that method is defined as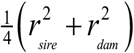, which leads to smaller values compared to those for 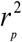 in **Table 1**. The estimated effective number of the candidate’s observations was always overestimated by the traditional method, and the same is true for the reliabilities of the dEBVs. That overestimation was only mild when the reliability was at a higher level, as in examples one and two, but more pronounced in the other examples with lower reliabilities. In particular, the old method did not recognize cases with a zero number of effective observations and zero reliability of the dEBV. A substantial overestimation was also observed for example three, where all information is contributed by offspring. Despite of overestimated 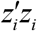 values in examples one, two, and four, the reliabilities of dEBVs were more similar to what was obtained by the new method. Also, the old method was not able to correctly replicate the number of effective observations 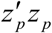 for the parent average in any of the examples. It should be noted that the reliability of the dEBV determines the weight that is used in the second stage of the analysis, while the size of 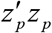 and 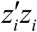 affect the size of the dEBV. This is why all three quantities are of importance in the deregression process.

**Figure 1** summarizes results from simulated data; scatter plots of traditional EBVs versus their newly derived counterparts show a strong linear relationship. Deviations from the diagonal dotted line in each panel indicate bias in the old method due to the relationship of the parents and the candidate’s inbreeding. The absolute size of the dEBVs was underestimated by almost exactly 10% in example one (**Figure 1**, left panel; linear regression coefficient 0.901, SE±0.002), where the reliability of dEBVs was considerably higher at 0.61 (Table 2) than in most other examples, with only a little difference between methods. In contrast, the reliability in example three was comparatively low at 0.12, and clearly overestimated in the old method. This was associated with a heavy 45% underestimation of the absolute size of the dEBVs when the the old method was applied (**Figure 1**, right panel; regression coefficient 0.550, SE±0.004).

**Figure 1.**
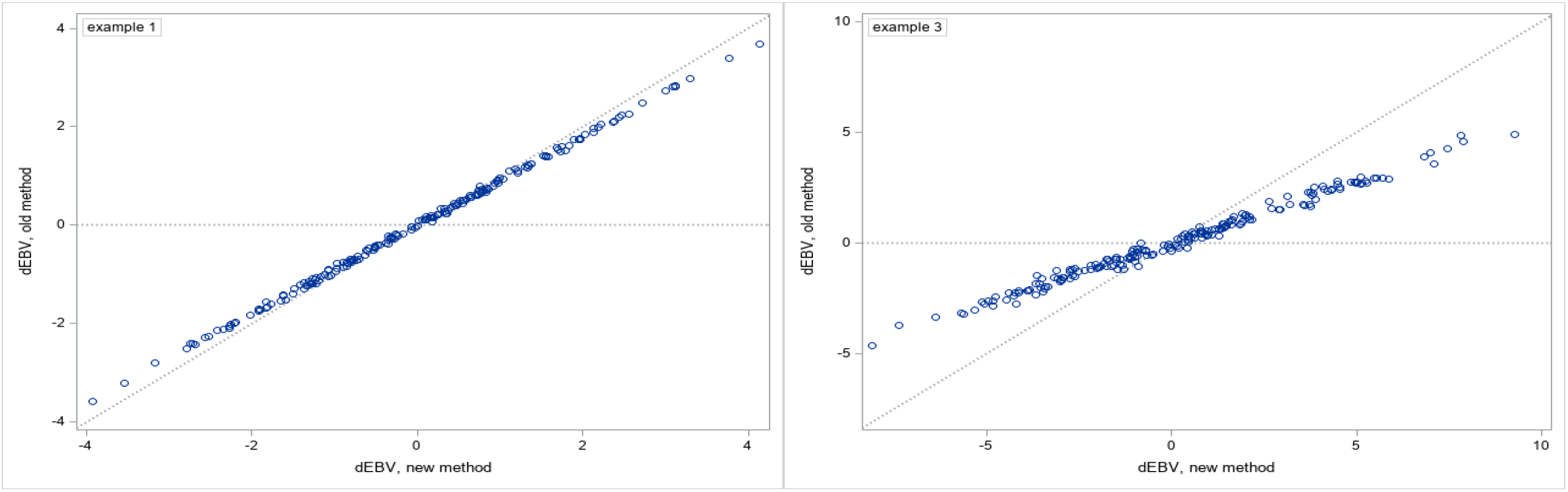
Scatter plots of dEBVs derived with the traditional method versus dEBVs from the new method. Each data point represents one repetition of 100 first-example (left) or third-example data (right) simulations. Deviations from the dotted diagonal line indicate bias in the old method, caused by inbreeding.

## Discussion

The examples show that, in the presence of inbreeding, the new method avoids the bias in the effective number of observations, the size of dEBVs, and their weights that is inherent in the traditional method. The improved method capitalizes on the three-by-three prediction error covariance matrix for each parent-candidate trio. In real applications, these matrices are not only shaped by relationships and inbreeding, but also by available information on other fixed and random factors, such as common litter environment or permanent environmental effects. By construction, the latter are excluded from the examples. Consequently, the observed bias can be fully attributed to inbreeding and relationships.

The second polynomial has coefficients{*β*_2_, *β*_1_, *β*_0_}that were given above as functions of reliabilities, among them the reliability of the candidate’s estimated Mendelian sampling term. As already mentioned, the dEBV is not defined in case this reliability is zero, and the weight is zero. Obviously, such observations have to be filtered out and cannot enter the second stage of analysis. Therefore, this reliability should be calculated from the three-by-three prediction error covariance matrix, no matter which set of coefficients is employed, and used as the criterion of choice for filtering out uninformative EBVs.

It is possible to express the coefficients of the second polynomial{*β*_2_, *β*_1_, *β*_0_}in terms of PEVs rather than reliabilities, and{*γ*_2_, *γ*_1_,*γ*_0_} in terms of reliabilities (data not shown), with no change in the resulting polynomials. In principle, both polynomials deliver identical dEBVs. However, their numerical stability differs: the second polynomial tends to be ill-conditioned against rounding errors in the PEVs. This was observed in an application to mouse data (Ding et al., 2026) and led to negative discriminants in the quadratic equation for a fraction of candidates, and to implausible reliabilities of the dEBVs of others (larger than the reliability of the EBVs). In contrast, the first polynomial performed well with the same data. The phenomenon could be reproduced with artificially created data (not shown) by rounding PEVs to a smaller number of digits. To prevent such difficulties, request more significant digits when retrieving PEVs with packages like ASReml. In general, the first polynomial is preferred because of its better numerical stability.

## Conclusions

The improved method relies on prediction error covariances for the EBVs of each candidate and its parents. It avoids biases in deregressed EBVs and their weights caused by inbreeding, especially when EBVs have a low reliability. Further, it enables consistent filtering of uninformative EBVs, preventing meaningless observations from being passed to the second stage of analysis.

## Appendix

The numerator-relationship matrix for the parents, the candidate, and the candidate’s two offspring was

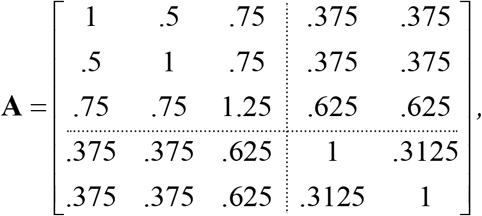

resulting in *a*_*pp*_ = 0.75 *a*_*mm*_ = 0.50 *a*_*bv*_ = 1.25 .

The parents are not inbred but related, with a relationship coefficient of 0.5. The ratio a_pp_/a_ii_ =1.5 deviates from one, due to the inbreeding of the candidate.

In example three, the candidate’s breeding value is related to one observation of each of its two progeny (animals 4 and 5), which both have no other known parent represented in the matrix. Thus, the expected value of 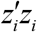 results from two weighted observations that are related with 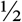 to the candidate’s BV. Thus, the diagonal element is 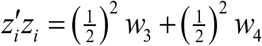, where *w* = *w* is the inverse ratio of the residual variance of an offspring of the candidate with the other parent unknown and the ordinary residual variance. There are three contributions to the first variance: the residual variance 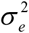, the genetic variance contributed by the unknown parent 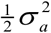, and the Mendelian sampling variance of the candidate’s gamete 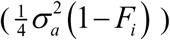. So,

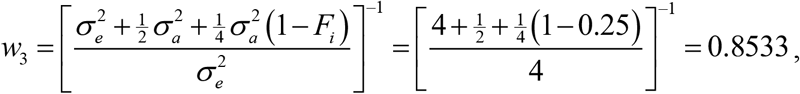

and the effective number of observations in example three is 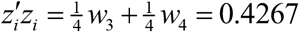.

Likewise, in example four, we get 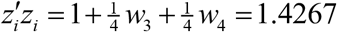, because the candidate has a phenotype plus two offspring.

## Notes

### Competing Interest Statement

The authors have declared no competing interest.

